# Higher levels of antibiotic resistance are less competitive: the hidden ecological cost of no-metabolic cost resistance

**DOI:** 10.64898/2025.12.15.694367

**Authors:** Miles T. Wetherington, Raymond Copeland, Christopher Zhang, Brian K. Hammer, Peter J. Yunker

## Abstract

Antibiotic resistance is often assumed to be constrained by fitness costs that limit the spread of highly resistant strains. Yet, many resistance mechanisms – including enzymatic antibiotic degradation – can arise with little or no metabolic cost, raising an important question: why is extreme resistance not more widespread? Here, we show that community-level interactions impose a hidden ecological cost on high resistance. By performing experiments with simple communities comprised of antibiotic resistant clinical isolates and an antibiotic susceptible strain, we find that when exposed to betalactam antibiotics, strains with a higher degree of antibiotic resistance can promote the survival of cohabiting susceptible strains. Guided by mean-field modeling, we find that highly resistant bacteria accelerate detoxification of the shared environment, shortening the period during which resistance confers a competitive advantage. Experiments with an engineered strain with tunable resistance level confirm that susceptible cells grow best in the presence of highly resistant strains. Importantly, this effect does not require evolved cooperation or active enzyme secretion; experimental and modeling results show that unavoidable processes, such as cell death or passive leakage, prevent complete privatization of resistance, giving rise to “accidental co-operation”. These findings suggest that resistance evolution is not only shaped by intrinsic cellular costs but also by ecological feedback that limits the benefits of incremental increases in resistance. This result may be reflected in the phenotypic responses of the clinical strains tested in this work, which fell into distinct low- and high-resistance classes with no intermediate phenotypes. Thus, this work demonstrates the important role of community dynamics in understanding the evolution of antibiotic resistance and treatment outcomes.

## Introduction

The discovery of antibiotics ushered in a new age of medicine, but the rise of antibiotic resistance threatens many of these advances. While antibiotic resistance is becoming more common, its spread is hypothesized to be limited as higher levels of antibiotic resistance incur proportionally greater costs [1–3], an idea consistent with experimental observations of a step-wise evolution of high levels of antibiotic resistance in the presence of increasingly higher concentrations of antibiotics [4, 5]. However, recent work has shown that increases in the level of antibiotic resistance can also occur with little-to-no metabolic cost[1, 6]. Bacteria can, for instance, boost antibiotic degradation by increasing enzyme production without significant metabolic penalties [1, 6], or through structural enzyme modifications that enhance catalytic efficiency independently of production costs [7–11]. A high level of antibiotic resistance with no metabolic cost would seem likely to rapidly sweep through a population; thus, if high, cost-free resistance is readily accessible, it is unclear why such highly resistant strains are not even more widespread in nature or the clinic.

In a related vein, while studies of antibiotic resistance typically focus on the responses of individual strains, a growing body of evidence suggests that community interactions are important. Bacteria are inherently social organisms, existing in dense, interconnected communities where interactions are frequent and significant—even within our own microbiome[12–15]. Crucially, many studies have demonstrated that susceptible bacteria can survive in the presence of resistant bacteria [16–24]. Such observations are often interpreted as microbial ‘public goods dilemmas.’ Such game theoretic scenarios are common for microbes, which perform many parts of their metabolism outside their bodies [25–27]. In these scenarios, ‘cooperators’ produce and secrete a public good while nearby ‘cheaters’ benefit from the presence of the public good without paying any cost to make it. Antibiotic resistance due to the production of antibiotic hydrolyzing enzymes would seem to map well onto such a game: resistant bacteria (cooperators) produce enzymes that degrade antibiotics, thus detoxifying their environment, while nearby susceptible bacteria (cheaters) reap the benefits of such enzymes without paying the cost of making them. In fact, Gram-negative bacteria make use of a classic solution to a public goods dilemma—they can retain the enzymes they produce within their periplasmic space, privatizing this otherwise public good [8, 10, 28–30]. This privatization is enabled by the fact that enzymes are typically ~three times larger than the outer membrane and peptidoglycan layer pore size [31]. Such enzyme retention has been recognized as a common mechanism to reduce cheating and stabilize cooperation in microbial populations [16]. Thus, it would appear that there are few barriers preventing high levels of antibiotic resistance in Gram-negative bacteria—increased effectiveness can be had at little-to-no cost, and the results can readily be privatized allowing the resistant strain to retain more of the benefit. In this context, it is unclear why high levels of antibiotic resistance are not even more prevalent.

Here we examine how antibiotic resistance levels shape the outcome of competitions between resistant and susceptible strains, focusing on beta-lactam antibiotics in simple two-strain communities across a range of drug concentrations. We first investigate the viability of susceptible *Vibrio cholerae* populations exposed to supra-MIC antibiotic concentrations when co-cultured with clinical isolates of *Acinetobacter baumannii*. These initial experiments motivate a minimal “toy model” that captures the key processes we expect to govern the system. Guided by this model, we experimentally dissect how varying resistance levels—controlled via inducer-driven beta-lactamase expression—affect the dynamics of co-cultured resistant and susceptible *V. cholerae* strains. Unexpectedly, we observe increasing susceptible fractions as resistance levels rise, contradicting the anticipated privatization of periplasmic beta-lactamase in Gram-negative bacteria. We demonstrate that this counterintuitive outcome arises from passive, non-growth-associated detoxification of the environment, rather than active growth of the resistant population *per se*. Returning to the *V. cholerae–A. baumannii* co-culture experiments, we find that (1) more resistant clinical isolates lead to higher final abundances of the susceptible strain, and (2) detoxification alone disproportionately benefits the susceptible strain when co-cultured with these highly resistant isolates. We discuss the implications of these findings for microbial community dynamics, antibiotic treatment outcomes, and the ecological consequences of resistance heterogeneity.

## Results

### Resistant Strains Rescue Susceptible Cells Under Antibiotic Stress

Motivated by these unresolved questions, we first measured how carbenicillin susceptible *V. cholerae* fare when co-cultured with resistant partners across a range of antibiotic concentrations. To do so, we co-cultured *V. cholerae* with 40 clinical isolates of *A. baumannii* at 0, 1, 2, and 4× the minimum inhibitory concentration (MIC) of carbenicillin for the susceptible *V. cholerae* strain (see Methods). For each resistant–susceptible pair, we quantified coexistence, lag time, and the final relative abundance of the susceptible population. Across many combinations, resistant strains were indeed able to rescue susceptible cells from extinction (*n* = 28). However, even in these cases we observed a consistent extension of susceptible lag times and a reduction in final susceptible abundance relative to matched co-cultures lacking antibiotics (Fig. 1A).

**Figure 1:**
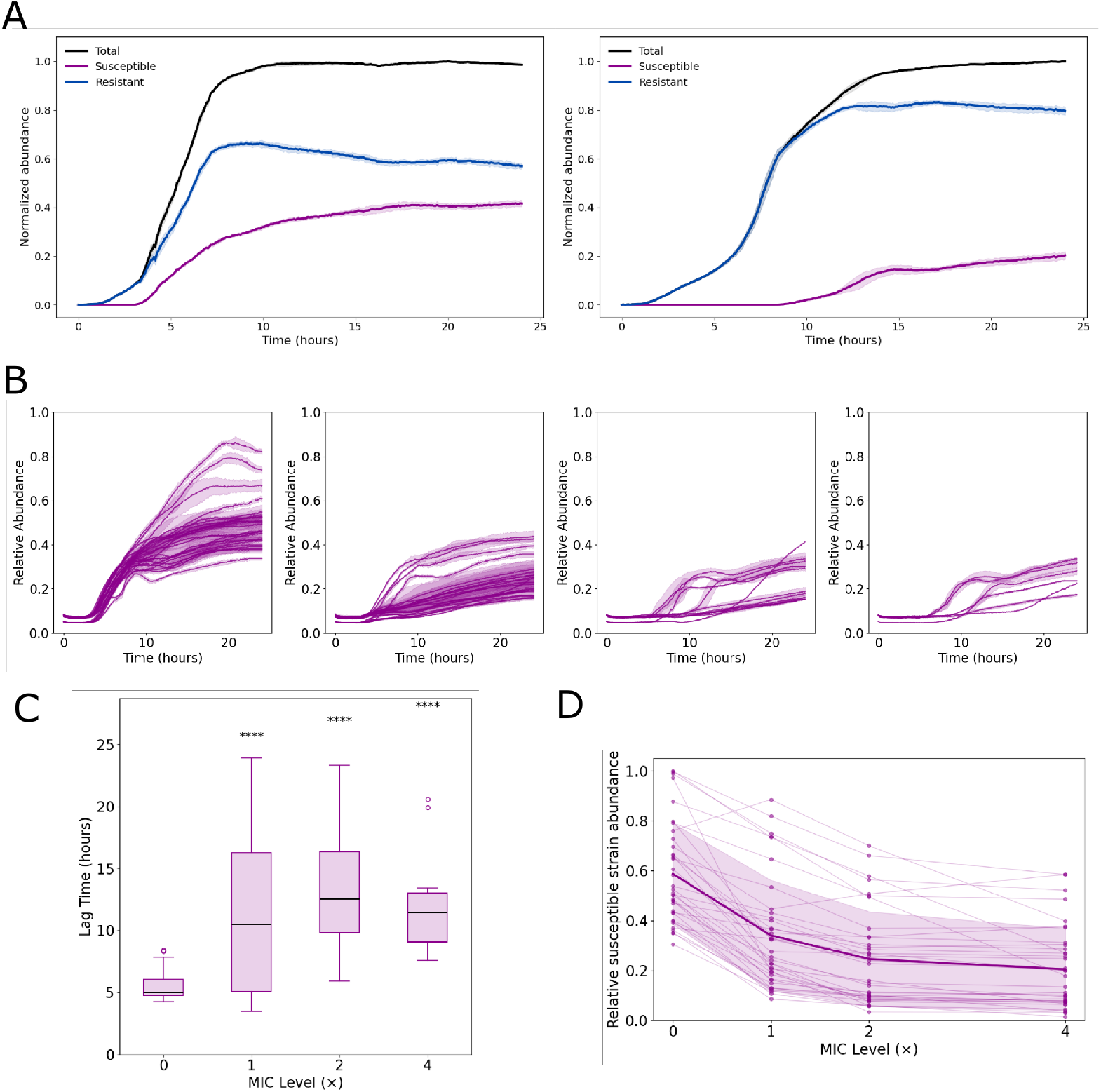
A. An example of the growth dynamics of susceptible strain - *V. cholerae* - (Magenta), resistant strain - *A. baumanii* - (Blue) and total (Black) under no antibiotics (left panel) and 4x MIC of susceptible strain (right panel) B. The relative abundance of *V. cholerae* grown with different isolates of *A. baumanii* at 0, 1, 2, 4x MIC C. The mean lag time of the growth curves of *V. cholerae* from B. Significant difference in mean lag time between 0x MIC and all antibiotic conditions for *V. cholerae* strain (*p <* 0.0001) D. The steady-state abundance of *V. cholerae* paired with 28 different clinical isolates of *A. baumanii* each pairing has *n* = 3 technical replicates.

Examining the susceptible population alone across all 40 co-cultures and all antibiotic concentrations, we found that only a subset of resistant–susceptible pairs permitted coexistence at the highest antibiotic concentrations (Fig. 1B). Overall, increasing antibiotic levels produced two consistent effects: (i) the relative abundance of the susceptible strain was reduced and (ii) susceptible strain lag times were longer and more variable. This pattern is readily visible when comparing susceptible lag times across the four antibiotic conditions in Figure 1B and quantified in Figure 1C. Furthermore, despite a broad range of differences in the competitive advantage between strains under antibiotic-free conditions, the relative abundance of the susceptible strain decreased across all coexisting pairs as antibiotic concentrations increased, as expected (Fig. 1D).

These results prompted us to seek a general mechanism that could explain both the extended lag times and the shifts in susceptible abundance. Given that beta-lactamase production is the predominant mode of beta-lactam resistance in Gram-negative bacteria, we next developed a minimal model capturing beta-lactamase activity and its impact on population dynamics.

### A Minimal Model of Beta-lactamase Activity and Population Dynamics

To understand what could cause these counterintuitive experimental observations, we developed a mathematical mean-field model exploring the broader implications of passive enzyme clearing on community dynamics. This model tracks the abundance of resistant and susceptible strains as fractional volumes of a fixed carrying capacity, along with internal and external concentrations of antibiotics and antibiotic resistant enzymes. It incorporates logistic growth constrained by available space, random cell death, antibiotic-induced death dependent on intracellular antibiotic concentration exceeding a threshold (the MIC), and enzyme-mediated antibiotic degradation, with diffusion of antibiotics in and out of the cells modeled by Fick’s first law. The model assumes resistant strains produce enzymes without growth rate costs. After cells die, their enzymes and antibiotics are released into the external environment to ensure mass conservation.

Using this model we look at time series data of the abundances of resistant and susceptible cells. First, we consider a simulation with a low level of enzyme efficacy (Fig. 2A). As expected the resistant cells grow while the susceptible cells die. In addition to following growth dynamics, we also track the dynamics of the external concentration of antibiotics (Fig. 2B). The time at which the external antibiotic concentration (green line Fig. 2B) drops below the MIC of susceptible (Horizontal dashed line Fig. 2B) indicates when the susceptible strain can start increasing abundance. When the enzyme efficacy is high (Fig. 2C), the environmental antibiotic concentration drops rapidly (Fig. 2D), reaching below the susceptible cell MIC before the abundance of susceptible cells has decreased by even a factor of 10. At this point susceptible abundance begins to increase. This “rebound effect” could explain why we experimentally observed the survival of susceptible cells—the resistant cells detoxify the environment. Extending this investigation we next explore the effect of enzyme efficacy of the resistant strain on outcome for susceptible populations at different initial abundances (Fig. 3A). Beyond a certain threshold, the effect of antibiotics in the environment appears to be negligible for susceptible strategies.

**Figure 2:**
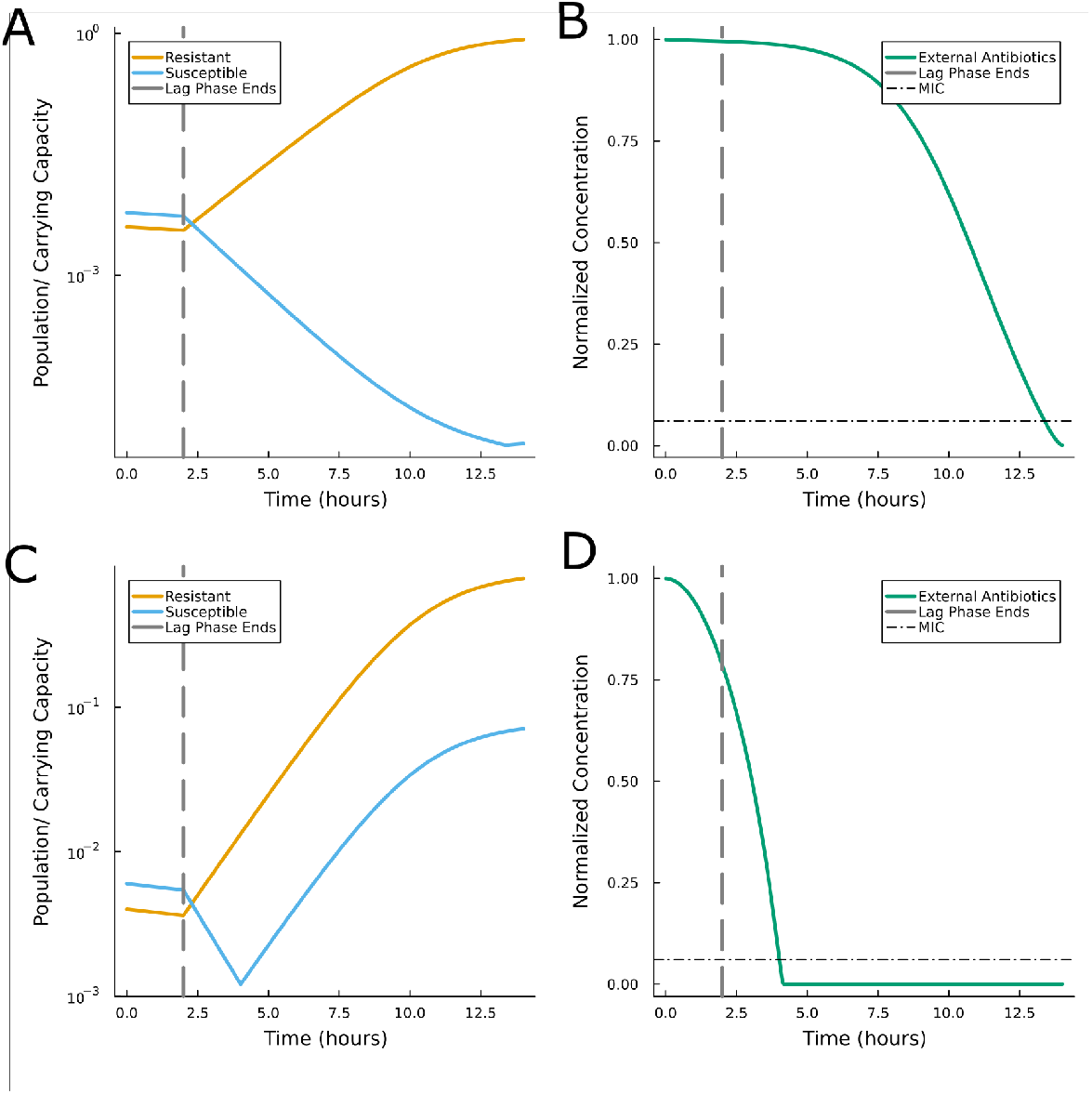
A. Growth dynamics of a 2-strain system with susceptible (blue) and resistant (orange) strategies. Dashed grey vertical line delineates end of lag-phase and beginning of log-phase for both strategies. Resistant has low enzyme efficacy. B. The dynamics of antibiotics degradation following A. horizontal dashed line is the minimum inhibitory concentration (MIC) for the susceptible strategy in A. Once concentration drops below this line the susceptible strategy can begin to grow C. Same as A. with high enzyme efficiency for the resistant strategy D. Same as C., high enzyme efficiency leads to quicker clearance of external antibiotics.

**Figure 3:**
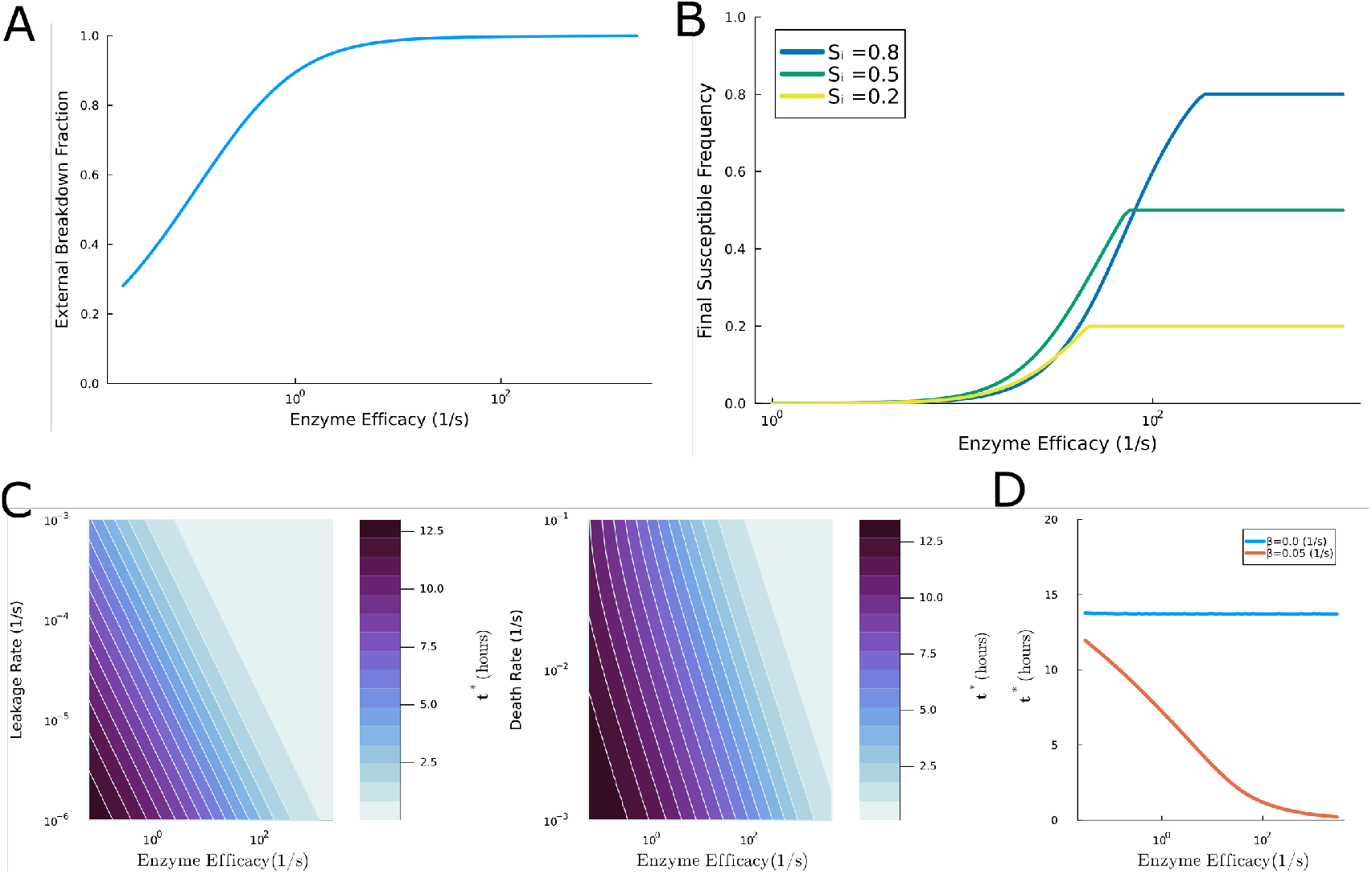
A. Fraction of enzyme breakdown occurring externally as a function of enzyme efficacy B. Final abundance of the susceptible strategy (y-axis) starting at different initial relative abundances and paired with resistance strategies with different levels of enzyme efficacy (x-axis) C. Passive leakage (left panel) and cell death (right panel) can create the conditions for “accidental cooperation” measured here by the change in the time until the external antibiotic concentration has dropped below susceptible MIC (*t*^*^) D. Comparison between *t*^*^ with (red) and without (blue) cell death.

To investigate what drove the breakdown of antibiotics facilitating the growth of susceptible cells we determined how much of the antibiotics were hydrolyzed inside resistant cells in our model. Surprisingly, we found that even for moderate efficacies (*k*_*cat*_ ≈ 10^1^(1*/s*)), less than 1% of antibiotics were broken down inside cells and over 99% of antibiotics were broken down outside cells (Fig. 3A). In contrast, for low efficacies (*k*_*cat*_ ≈ 10^−2^(1*/s*)), 67% of antibiotics were broken down inside cells and and 33% of antibiotics were broken down outside cells. Therefore, we find that as efficacy increases external antibiotic break down becomes the dominant process. This result is surprising as there is no enzyme secretion or leakage in our model. Thus, random cell death alone must be sufficient to facilitate community-wide antibiotic detoxification. To test this, we varied the enzyme efficacy and random death rate and tracked the time it took for the environmental antibiotic concentration to break down below the MIC (Fig. 3C, right panel); we refer to this time as *t*^*^. We find that *t*^*^ rapidly decreases as enzyme efficacy or the death rate increases. Taking these assumptions at face value it is clear: within this simple model, resistant populations experience an inevitable “accidental cooperation” in co-culture scenarios.

We next sought to determine if enzyme leakage could be a plausible alternate route to cooperation by accident (Fig. 3b, left panel). A diffusion based uptake and leakage term was added to our system (see equation 10) with the same form as the antibiotic uptake and leakage term where the flux between the internal and external environments is based on the volume of the cells, the difference in concentration, and a set leakage rate. We find that even a slow enzyme leakage rate (10^−4^*/s*, meaning a single cell would take roughly three hours for leakage to equilibrate its internal environment to its external environment), can lead to a small *t*^*^. What is abundantly clear from these model assumptions is that one of these mechanisms must be present to see the effects presented here. We performed simulations without random death and without enzyme leakage and measured *t*^*^. Absent ‘cooperation by accident’ mechanisms, the time for antibiotic breakdown is long and relatively insensitive to enzyme efficacy once a strain is resistant (Fig. 3D). This is because in order for a strain to be resistant, it must be able to breakdown most of the incoming antibiotics. Interestingly, diffusion of the antibiotics inside the cell becomes the rate limiting step not the hydrolysis rate, so the breakdown time does not change substantially with increasing efficacy. Therefore, full privatization of beta-lactamase enzymes appears inherently impossible in realistic biological contexts. We thus hypothesize that “accidental cooperation” is responsible for our experimental observation that strains with higher levels of antibiotic resistance perform worse than resistant strains with lower levels of antibiotic resistance in co-culture with susceptible strains despite no growth rate cost.

### Increased Resistance Elevates Susceptible Abundance via Antibiotic Degradation

Next, to systematically investigate how enzyme production affects the dynamics of antibiotic resistance within microbial communities we engineered and experimentally characterized a resistant version of our *Vibrio cholerae* strain to express beta-lactamase enzymes at levels controlled by an inducer (see Methods for details on the strain). We verified that increased inducer concentrations increased the level of antibiotic resistance against carbenicillin. First, we checked to confirm that enzyme production did not impose significant metabolic costs. We cultured the resistant strain with a range of antibiotic concentrations and inducer strengths, i.e., resistance levels (Fig. 4A). Indeed, increasing the inducer concentration without the presence of antibiotics does not lead to slower growth (See first column of Fig. 4A) suggesting a negligible cost to betalactamase production. With the presence of antibiotics - increasing the inducer concentration for a fixed antibiotic concentration led to a decreased lag time (Fig. 4B, left panel). Conversely, increasing the antibiotic concentration for a fixed inducer concentration extended the lagtime (Fig. 4B, right panel). Both of these observations are consistent with the assumptions built into our model.

**Figure 4:**
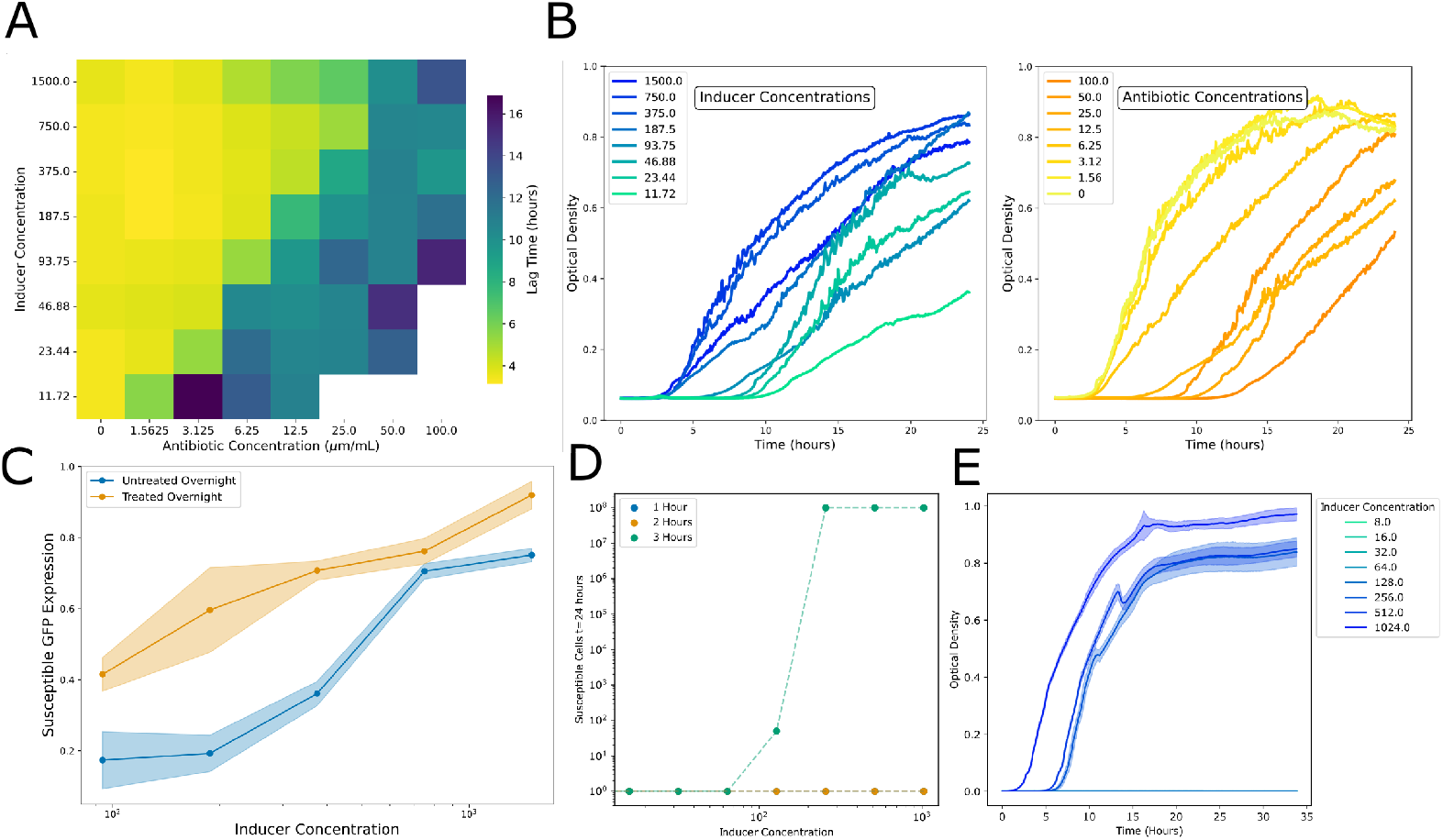
A. Lag time for our engineered resistant strain of *V. cholerae* under different concentrations of carbenicillin and inducer (theophylline). Empty region indicates extinct threshold B. Example growth curves from cross sections along A. along inducer concentrations (left panel) and antibiotic concetration (right panel) C. Relative susceptible *V. cholerae* abundance when grown in co-culture with resistant *V. cholerae* under 4x MIC and different inducer concentrations of inducer (*n* = 5). Treated (untreated) had corresponding inducer concentration included (excluded) in overnight D. Antibiotic degradation leakage assay: resistant *V. cholerae* at 7 different inducer concentrations (x-axis) is grown for *t* = 1, 2, 3*h* with 4x MIC of carbenicillin for susceptible strain before removed and media is inoculated with susceptible *V. cholerae*. Final abundance after 24 hours (y-axis) E. Antibiotic degradation cell death assay: resistant strain is grown with different inducer concentrations overnight, removed from conditioned media and immediately 4x MIC carbenicillin and susceptible cells are added then grown for over 24h.

We next considered how resistance level influences community composition by co-culturing susceptible and resistant populations at equal initial proportions (Fig. 4C). Interestingly, increasing the resistance level (inducer concentration) led to a higher final proportion of susceptible cells. These results are consistent with predictions from the model. We thus next sought to determine if the accidental cooperation that drove the phenomenon in our model is also present in these experiments.

To test the accidental cooperation hypothesis experimentally, we performed sequential inoculation assays. Resistant cells expressing varying enzyme levels (i.e., in the presence of different inducer concentrations) grew alone in antibiotic environments for different durations before being removed leaving behind the conditioned media (Fig. 4D). Subsequently, susceptible cells were introduced into this conditioned environment. Remarkably, we found that susceptible populations successfully grew only after resistant cells had been allowed to proliferate for at least three hours at sufficiently high resistance levels. This experiment demonstrates that enzyme-mediated detoxification of the antibiotic environment occurs passively, without requiring direct interaction or coexistence between strains. As an additional test, we took overnight cultures of the resistant strain – grown without antibiotics – and filtered the cells out. We then added antibiotics and susceptible cells simultaneously. We found that susceptible cells were able to grow in environments that had contained resistant cells at the three highest inducer concentrations conferring the highest resistant levels (Fig. 4E). Together, these observations demonstrate that the presence of resistant cells is not necessary to condition the media; instead, the beta-lactamase the resistant population leaves behind is sufficient to detoxify the environment and thus enable the growth of susceptible cells. Such passive detoxification thus provides indirect protection to susceptible cells and helps explain the observed rise in susceptible proportions.

After verifying these results with this simple lab 2-strain system where we ensured that betalactamase production was the source of resistance – and in turn accidental cooperation – we returned to our *V. cholerae* - *A. baumanii* community. Our first task was to determine the resistance levels of our clinical strains. To our surprise resistance levels among clinical isolates permitting susceptible growth fell neatly into 2 discontinuous levels of low (≤ 64*µg/mL*) and high (≥ 512*µg/mL*) resistance (Fig. 5A). These two categories broadly showed different behaviors when in co-culture with our susceptible strain. Indeed, as expected from the previous results we find that high resistant clinical strains facilitated greater survival than low resistant clinical strains (Fig. 5B). For example, after co-culture at 4X the susceptible *V. cholerae* MIC, the relative abundance of susceptible cells was 0.288 *±* 0.184 and 0.077 *±* 0.131 (p-value = 1.607e-3) for high resistance and low resistance clinical isolates, respectively (Fig. 5C). Crucially, this phenomenon occurs despite the fact that there was no significant difference in competitive ability between low and high resistance strains under antibiotic-free conditions with the susceptible strain; specifically, the mean relative abundances of the susceptible strain were 0.632 *±* 0.219 and 0.576 *±* 0.213 when co-cultured with the low and high resistance strains (p-value = 0.509; Fig. 5C).

**Figure 5:**
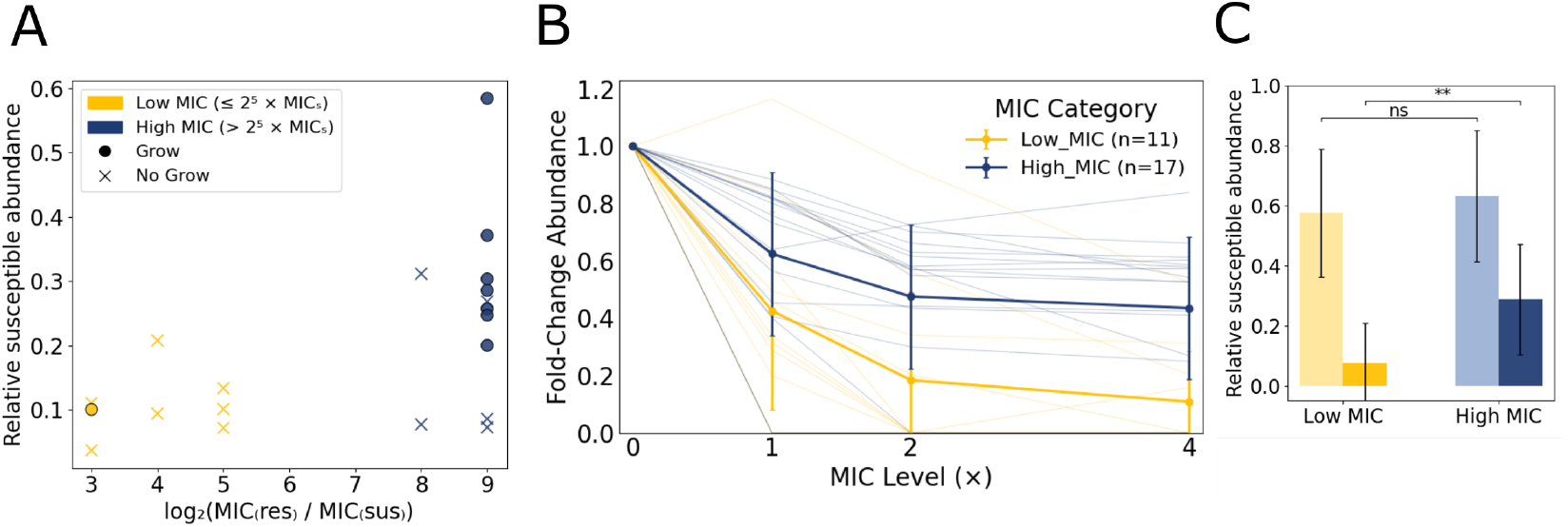
A. The relative abundance of susceptible *V. cholerae* (y - axis) for all pairs with resistant clinical strains *A. baumanii* with respect to level of relative resistance (x-axis). Resistant clinical strains fall into 2 natural categories of low (yellow) and high (blue) separated by unoccupied intermediate resistance region: high (MIC ≥256x MIC Susceptible, *n* = 17) or low (MIC ≤ 32x MIC Susceptible, *n* = 11). Susceptible strains that grew (did not grow) in conditioned media in growth assay similar to Fig4E indicated with circle (x). B. Fold-change abundance of susceptible *V. cholerae* relative to 0x MIC in co-culture with different clinical strains of resistant *A. baumanii* categorized as high or low. Average (bold line) for low (yellow) and high (blue) resistant strain pairing. C. Mean susceptible *V. cholerae* abundance in no antibiotics (left bar in each pair) and 4x MIC of susceptible (right bar in each pair) for low (yellow) and high (blue) resistant category. No significant difference between relative growth of susceptible strain when grown with either high or low resistant clinical *A. baumanii* strain; 0.632 *±* 0.219 and 0.576 *±* 0.213 (p-value = 0.509). Significantly higher abundance of susceptible when grown in co-culture with high resistant strain in 4x MIC antibiotics than when grown with low resistant strain; 0.288 *±* 0.184 and 0.077 *±* 0.131 (p-value = 1.607e-3).

Finally, we repeated the accidental cooperation experiments with the clinical strains. Overnight cultures were produced, and the cells were removed. Subsequently, antibiotics and susceptible populations were inoculated. Here we specifically track where susceptible populations did and did not grow with respect to the resistance level of the strain that conditioned the media (Fig. 5A). This result shows a clear trend that higher resistance confers greater environmental detoxification, supporting the idea that cooperation by accident facilitates the effects we observe.

## Discussion

Together, our results demonstrate that increasing levels of antibiotic resistance can promote the survival of susceptible strains in mixed microbial communities. Across both clinical isolates and engineered strains, resistant partners rescued susceptible populations under otherwise lethal antibiotic conditions, but did so in a manner accompanied by extended lag phases and reduced final abundances. A minimal mechanistic model shows that these dynamics emerge naturally from beta-lactamase-mediated antibiotic degradation acting on the shared environment, and that sufficiently high resistance accelerates detoxification in ways that disproportionately benefit susceptible neighbors. Critically, both modeling and experiments indicate that this effect does not require evolved cooperation or direct interaction between strains: passive processes such as random cell death or minimal enzyme leakage are sufficient to prevent full privatization of resistance. This passive sharing mechanism represents a unique case of “accidental cooperation,” a form of unavoidable mutualism in microbial communities where benefits to other community members emerge not through intentional secretion but as an inevitable consequence of cell mortality or passive leakage [32]. In classic public goods scenarios, privatization strategies typically limit exploitation by cheaters, yet we demonstrate that in beta-lactamase-mediated resistance, privatization is ultimately temporary, constrained by biological realities such as random cell death and lysis.

What makes these dynamics particularly counterintuitive is that increasing enzyme efficacy can actually diminish the fitness advantage of resistant bacteria rather than enhancing it, without a growth rate cost. This occurs through two interlinked mechanisms. First, higher enzyme efficacy accelerates environmental detoxification, shortening the critical period during which antibiotic presence confers a selective advantage to resistant strains. Second, resistant strains face a “ceiling effect”: as the good they produce does not enhance their growth rate, but instead decreases their death rate. Thus, the rate at which they expand their population has a set maximum independent of the enzyme efficacy. As antibiotics are rapidly neutralized, this advantage diminishes as well. These mechanisms thus produce not just diminishing but decreasing returns on enzyme improvement, suggesting that community dynamics may play an important role in the evolution and maintenance of antibiotic resistance. That the good produced decreases death rate rather than increases growth rate is essential for these counterintuitive dynamics. As antibiotic death rates cannot be lower than zero, any additional efficacy beyond what is necessary for a single resistant cell to have an antibiotic death rate of zero cannot benefit the resistant cells, and thus can only benefit the susceptible cells. Consequently, selection may favor low, “good enough” enzyme efficacies, rather than incrementally more effective ones. This may help to explain the observation that clinical strains fall neatly into 2 distinct resistance strategies – “good enough” & highly resistant – with no intermediate. It would be interesting to survey a larger number of stains than the 28 studied here to see if such groupings persist.

In many cases the resistant strain may not benefit from the extinction of the susceptible strain. For example, the two strains may cooperate via cross-feeding [33, 34]. In such cases, accidental cooperation may represent a case in which a beneficial trait emerges “for free” i.e., without the resistant strain having to evolve mechanisms to secrete large enzymes. Instead, cooperation emerges via the biophysical breakdown of cell membranes after death, reminiscent of other biological traits that emerge for free from unavoidable physics [35–42].

Crucially, we see accidental cooperation emerge at efficacies that are, in fact, well within the empirically measured range. In our simulations, we see the susceptible strain’s abundance increases substantially at enzyme efficacies ~10^1^*/s*) (Fig. 3B), which is much lower than most beta-lactamase efficacy measurements found in the literature which can reach over 10^3^*/s* [7, 9–11, 43]. Further, our engineered strain produces cooperation by accident at promoter concentrations that correspond to MIC values ~200*µg/mL*, which is again within range of various strains resistant to carbenicillin [44]. Thus, this mechanism is unlikely to be one viewable only in artificial laboratory and in silico environments, but instead likely happens in nature and in many infections – a position strengthened by our results with clinical isolates.

The potential for the broad prevalence of cooperation by accident in antibiotic resistance challenges traditional approaches to predicting and managing antibiotic resistance in clinical and environmental contexts. Current methodologies evaluate resistance at the single-cell or single-strain level, potentially missing important community-scale dynamics arising from accidental cooperation. In fact, antibiotic treatment fails 10% of the time when susceptibility testing suggests a drug should eradicate the pathogen [45]. Further, for chronic infections, single strain antibiotic susceptibility testing is often not used in the selection of treatment plans due to its weak correlation with patient outcomes [46]. While there are many reasons for these treatment failures, cooperation by accident may be a contributing factor. Thus, incorporating these ecological interactions into resistance assessments could yield more accurate predictions and inform strategies designed to minimize unintended community-wide detoxification, thus preserving antibiotic efficacy.

## Methods

### Mean Field Population Model

To investigate multi-strain community dynamics under antibiotic pressure, we developed a mean field model tracking concentrations of antibiotics and enzymes inside and outside cells, influenced by previous works including [42] and [17]. We model *m* microbial populations (strain *i*, where *i* = 1 to *m*) with volumes *V*_*i*_, tracking these as proportions of carrying capacity (*K*) such that *N*_*i*_ = *V*_*i*_*/K* represents the fraction of maximum volume occupied by strain *i*. Remaining space is potentially occupied by cells is:

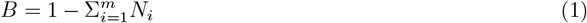

We constrain the total system volume (∑ _*i*_ *V*_*i*_ + *V*_*E*_ = *V*_Total_), where *V*_*E*_ is external volume. Population dynamics are governed by:

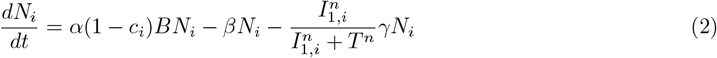

These terms represent logistic growth, random death, and antibiotic death, respectively. We implement a two-hour lag time at simulation start during which no reproduction or antibiotic death occurs, reflecting decreased metabolism when bacteria enter new environments [17, 47].

Random death occurs at rate *β*, an important function in microbial communities [48–50]. Antibiotic death follows two principles: antibiotics affect cells only when intracellular concentration (*I*_1,*i*_) exceeds threshold *T*, and decay rate is proportional to reproduction rate [51–53].

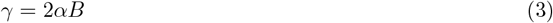

Here, subscript 1 refers to antibiotics and subscript 2 to enzymes. When antibiotic concentration exceeds threshold *T*, controlled by 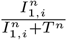, the *γ* term inverts logistic growth, causing decay proportional to potential growth rate. This models the observation that beta-lactams kill during reproduction [52, 54, 55].

For antibiotic dynamics, we use Fick’s first law of diffusion [43, 56]:

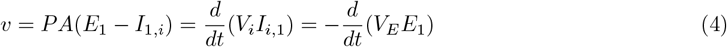

Replacing *P* with characteristic leakage rate *r*_*l*_ and *A* with cell volume:

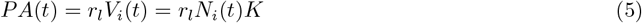

The resulting dynamics equations are:

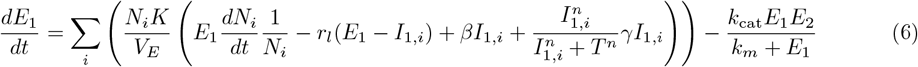

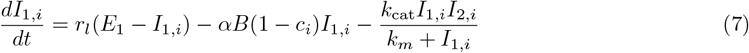

### Enzyme Dynamics

Our system has two strains: resistant (*R*) and susceptible (*S*). The *R* strain produces enzymes with distinct kinetic parameters, but there is no growth cost for enzyme production [6]. Enzymes exist both internally and externally due to passive diffusion and post-lysis release.

We assume resistant cells maintain constant internal enzyme concentration 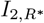:

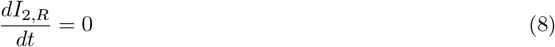

Susceptible strain enzyme dynamics:

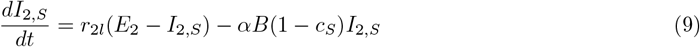

External enzyme dynamics:

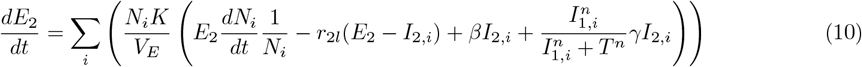

To begin, we will set leakage rate *r*_2*l*_ at zero to maximize enzyme privatization within producing strains, but later we will relax this.

### Single Cell Resistance

We can determine the minimum enzyme efficacy for single-cell survival at a given antibiotic concentration. Setting 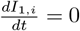 and assuming internal concentration equals threshold *T* :

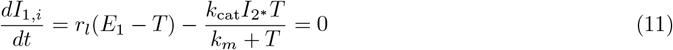

Solving for *k*_cat_:

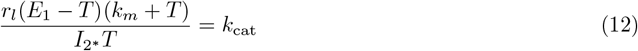

The minimum inhibitory concentration (MIC, *C*_0_), substituting *V*_*max*_ for *k*_*cat*_*I*_2_:

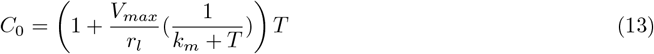

This aligns with previous analytical MIC predictions [43]. We define enzyme efficacy as a proportion of that required for single-cell survival at a given antibiotic concentration.

### Strain Construction

Our two *V. cholerae* background strains were derived from Thelin and Taylor [57] (designated BH1514) and Crisan et al. [58] (designated SN440). We used the Gibson Assembly method on Q5 polymerase PCR amplified DNA fragments to add one of two DNA constructs into a pEVS141 plasmid background (obtained from Waters et al. [59]). pCZ1 contains a Ptac-theophylline riboswitch inducer region followed by bla, an beta-lactamase antibiotic resistance gene, placed on the reverse strand translationally linked to mTFP1, a fluorescent protein. pCZ1 containing strains were experimentally shown to not produce any antibiotic resistance beta-lactamases proteins even with IPTG and theophylline induction. These strains were designated as susceptible strains in this study. pCZ2 contains a Ptac-theophylline riboswitch inducer region followed by bla translationally linked to mTFP1. Strains containing pCZ2 were designated as Resistant strains in this study. pCZ2 containing strains were experimentally shown to produce antibiotic resistance beta-lactamases proteins in an IPTG or theophylline dose dependent manner. Plasmid maps of these plasmids used in this study are available in the supplementary material. Recombinant plasmids were introduced into *V. cholerae* with the following method. Our plasmids were first transformed into S17 dap- *E. coli* obtained from Simon et al. [60] via electroporation. The resulting *E. coli* conjugation donor strain containing our s of interest were biparentally mated with our *V. cholerae* background strains using the method described in Burns and DiChristina [61]. The Ptac-riboswitch inducer region in the plasmids was obtained from Dalia et al. [62].

### Growth conditions

All experiments were performed in Muller Hinton (MH) Broth. Clinical isolates of *A. baumanii* used in this work has been previously published [63]. Antibiotics and inducer stocks were stored at −20C^°^. Prior to wellplate experiments, strains were grown straight from −80C^°^ stocks overnight for 16h before being back-diluted 1:1000 into 200*µ*L of MH in wells containing the necessary antibiotics and inducer concentrations. For co-culture wellplate experiments the constitutively produced green fluorescence by our *V. cholerae* strain was used as a proxy for susceptible biomass along with OD600 for total biomass allowing us to determine relative abundance of each strain in co-culture growth. Expression of GFP was normalized by a maximum value produced (Susceptible abundance = 1) when grown in isolation.

### Experimental Growth Curve Fitting

The growth rate (*α*) and lag times (*t*_lag_) were calculated by fitting the optical density time series to the following equation:

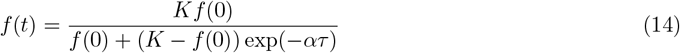

Where *f* (0) is the time-zero optical density, *K* is a fit parameter for the carrying capacity, *α* is the fit parameter for growth rate, and *τ* is defined as *τ* = max(0, *t* − *t*_shift_) and *t*_shift_ is the third fit parameter. The purpose of *τ* is to allow the curves to spend time *t*_shift_ with a zero growth rate before increasing at rate *α*. Note this also means after time *t*_shift_ the population increases as if this were time 0.

To define lag time, we first utilize the formula from [64] which defines lag time as the maximum curvature of logistic growth using the approximation:

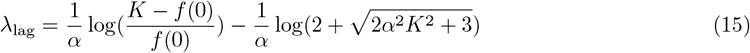

But this assumes the population begins growing at time-zero, so our reported lag times are the sum of these times:

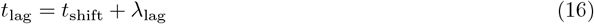

## Acknowledgments

This work was supported by NIH Grant No. 1R35GM138354-01 to P.J.Y.

